# Circulating plasma fibronectin affects normal adipose tissue insulin sensitivity and adipocyte differentiation

**DOI:** 10.1101/2024.02.28.582553

**Authors:** Mahdokht Mahmoodi, Elahe Mirzarazi Dahagi, Mir-Hamed Nabavi, Ylauna Penalva, Amrita Gosaine, Monzur Murshed, Sandrine Couldwell, Lisa Munter, Mari T. Kaartinen

## Abstract

Plasma fibronectin (pFN), a liver-derived, circulating protein, has been shown to affect adipocyte morphology, adipogenesis, and insulin signalling in preadipocytes *in vitro*. In this study, we show via injections of fluorescence-labelled pFN to mice *in vivo* its abundant accrual visceral and subcutaneous adipose tissues (VAT and SAT). Diet-induced obesity model of liver-specific conditional *Fn1* knockout (pFN KO), showed no altered weight gain or differences, whole-body fat mass or SAT or VAT volumes after 20- week HFD-feeding, however, mice showed significantly improved glucose clearance and whole-body insulin sensitivity on normal diet. Furthermore, *in vivo* insulin sensitivity assay revealed significant increase in AKT phosphorylation in pFN KO SAT on normal diet as well as in normal and obese VAT of the pFN KO. Histological assessment of the pFN KO depots showed significant increase in small adipocytes on normal diet, which was particularly prominent in SAT. RNA sequencing of the normal diet-fed pFN versus control SAT revealed alterations in fatty acid metabolism and thermogenesis suggesting presence of beige adipocytes. VAT RNA sequencing after HFD showed alternations in genes reflecting stem cell populations. Our data suggests that the absence of pFN alters cell pools in AT favoring cells with increased insulin sensitivity.

## 1. Introduction

Adipose tissue (AT) and adipocytes play an essential role in maintenance of whole-body energy balance and nutritional homeostasis, via acting as energy storage for both fat and glucose, and via balancing lipogenesis and lipolysis (1). AT expansion and lipogenesis occurs in energy surplus via processes involving adipocyte proliferation (hyperplasia), adipogenesis (adipocyte differentiation) and adipocyte enlargement (hyperptophy) all of which are regulated by insulin (1, 2). White adipose tissue (WAT) which represents energy storage depot, is classified as subcutaneous (SAT) or visceral (VAT)(3, 4). Brown adipose tissue (BAT) and beige fat, which can convert from WAT under specific conditions, are thermogenic with high density of mitochondria and ability to process fatty acids into heat (5). While both SAT and VAT depots expand in obesity, an increase in VAT (around internal organs), is more detrimental to metabolic health and correlates with development of whole-body metabolic dysfunction (4, 6). VAT adipocytes are larger and metabolically more active (more prone to lipogenesis/lipolysis) whereas SAT is more cellularized, more vascularized, less prone to inflammation in obesity (6, 7) and has capacity to convert to beige fat with increased thermogenesis (5).

Adipocyte lineage cells are insulin sensitive where preadipocytes, smaller adipocytes, and beige adipocytes are more insulin responsive compared to enlarged hypertrophic adipocytes (2, 7, 8). Whole-body and AT insulin sensitivity and signalling are physiologically modulated by plethora of factors including circulating myokines, adipokines, and cytokines (9–12). Locally, in AT, adipocyte insulin sensitivity and signaling can be regulated by the cellular microenvironment and extracellular matrix (ECM) that provides appropriate, lineage stage-specific adhesion-modes and stiffness-mediated signals to boost insulin signaling (13, 14). Intracellular insulin signals culminate to AKT (protein kinase B) phosphorylation which leads to glucose transport and uptake to cells (9, 15).

Fibronectin is a multifunctional ECM glycoprotein encoded by *Fn1* gene which is expressed by multiple cell types and as several isoforms (16–19). Two physiological pools of FN exist: circulating plasma FN (pFN) and cellular FN that is made by tissue-resident cells (20, 21). pFN is synthesized by hepatocytes and secreted into the bloodstream where it circulates at a high concentration of 200-600 μg/ml in humans, and 100-400 μg/ml in mice (17). pFN does not contain any extra domains that are frequently detected in cellular FN (cFN) (22, 23). Previous research indicates that circulating pFN can absorb into tissues and form ECM fibrils and contribute to tissue integrity (24, 25). Experiments where pFN has been labelled and injected to rodent models or genetically deleted has demonstrated that pFN integrate to and/or have roles in skin, kidney, liver, heart muscle, bone, lung, vasculature and brain (24–27) where it may form 90% or more of the total FN matrix (28). It has not been reported if pFN contributes to AT ECM *in vivo*, however, the important role of FN in general in adipocyte biology has been demonstrated via *in vitro* studies showing that FN ECM sustains preadipocyte phenotype and inhibits adipose conversion jointly with Preadipocyte factor-1(Pref1) (29–34). Our work has also shown that pFN, from cell culture serum, readily forms ECM and promotes preadipocyte pre-adipocyte proliferation and its decrease allows for pro-adipogenic insulin signaling (35). Furthermore, we showed that pFN assembly in adipocyte cultures is promoted by transglutaminase (TG) activity from plasma TG, Factor XIII-A (FXIII-A) which in turn is produced by pre-adipocytes; followed by decreased expression as preadipocytes differentiate, mature and accumulate lipid (35). FXIII-A null AT has both increased insulin sensitivity and decreased ECM, including FN matrix (36). The role of pFN *in vivo* in whole-body metabolism or in AT function has not been explored. In this study we have investigated the potential contribution of circulating pFN to WAT and metabolism *in vivo*. We report that pFN adsorbs from circulation to SAT, VAT and BAT and its elimination of pFN from circulation in a mouse model with a liver specific *Fn1* conditional knockout (pFN KO) enhances glucose clearance, whole-body insulin sensitivity, AT cellularity, and insulin sensitivity in mice on a normal diet. The phenotype is particularly evident in SAT where we also detected altered adipocyte pools including evidence of beige adipocytes. Development of high fat diet (HFD)-induced obesity in pFN KO mice revealed no change in weight or fat depot distribution with simultaneous disappearance of insulin sensitivity and enhanced glucose tolerance, however, VAT remained significantly more insulin sensitive compared to control and showed altered pathways referring to stem cell populations. Our work assigns a novel function to pFN in AT homeostasis and adipose tissue cell pools.

## 2. Methods

### Reagents and antibodies

Antibodies against pAKT (Ser473) pAKT (Thr308) and AKT (pan) (all rabbit polyclonal) and antibody against β-actin was from Cell Signaling Technology (Danvers, MA, USA). Antibody against fibronectin (ab6328, mouse monoclonal) was from Abcam (Cambridge, UK) and FXIII-A antibody (SAF13A-AP, sheep polyclonal) from Affinity Biologicals (Ancaster, ON, Canada). Dulbecco’s Modified Eagle Medium (DMEM), GlutaMAX (12561–056), penicillin-streptomycin, and sodium pyruvate were purchased from Gibco (Burlington, ON, Canada). Fetal bovine serum was obtained from Hyclone (Waltham, MA, USA). All other reagents if not defined were purchased from Sigma-Aldrich.

### Animals

C57/bl6 male mice (for plasma FN injections) (were purchased from The Jackson Laboratory (Bar Harbor, ME, USA). Liver-specific deletion of *Fn1* gene results in elimination of FN from circulation. Plasma FN knockout model (C57/bl6 background) was generated as before from breeding *Fn1*flx/flx and ALB-Cre driver mouse. The *Fn1* flx/flx mice, originally developed by Sakai et al., (37) were generously provided to us by Dr. Faessler from Max Plank Institute for Biochemistry (Munich, Germany) via Dr. Reinhardt from McGill University (Montreal, Canada) (model also available at Jackson Laboratory (JAX #029624). ALB-Cre mouse (JAX #003574) (38) was purchased from The Jackson Laboratory (Bar Harbor, ME, USA). All animals were housed in a pathogen-free animal facility and kept on a diurnal cycle and had ad libitum access to food and water. The animal care and experimental procedures were under the Canadian Council of Animal Care and approved by the McGill University Animal Care Committee.

### Plasma fibronectin labeling and detection in vitro and in vivo

pFN (human plasma FN (FC010) (Millipore,Burlington, MA, USA) was labelled with AlexaFluor568 or AlexaFluro680 according to instructions of the manufacturer (ThermoFisher, Waltham, MA, USA). Briefly, pFN was concentrated to 1 mg/ml solution with Amicon® Ultra 3K Centrifugal Filter Devices (3,000 molecular weight cut off) (Sigma-Aldrich, St Louis, MI, USA), followed by two sequential dialysis processes using dialysis cassettes (Slide-A-Lyzer™ Dialysis Cassettes, 10K MWCO) (ThermoFisher, Waltham, MA, USA) against 100mM NaHCO_3_ + 500mM NaCl pH 8.4 buffer. Protein concentration was measured using Micro BCA Protein Assay Kit (ThermoFisher, Waltham, MA USA). Dissolved AlexaFluor 568 NHS Ester (Invitrogen, Waltham, MA USA) in dimethyl sulfoxide (DMSO) at concentration of 10 mg/ml was added to the pFN solution, and the solution was incubated at room temperature (RT) for 1 h with constant and gentle shaking. The unbound dye was removed via two dialysis steps (as above) at 4°C against 100 mM NaHCO3 + 500mM NaCl, pH 8.4. BCA assay was used to determine the final protein concentration of AlexaFluor 568-pFN. Labeling degree was assessed according to the manufacturer’s recommendations and determined to be 3.5 AF568 molecules per one pFN molecule. The ability of pFN to create a fibrillar matrix, was confirmed in 3T3-L1 cells via adding AlexaFluor568-pFN or AlexFluro680 to media at 2 μg/ml concentration to (see below).

### 3T3-L1 cultures

3T3-L1 Mouse embryonic fibroblasts (MEFs)(ATCC, State, USA)) cells were cultured in 8-well Nunc Lab-Tek® II glass chamber slides (Fisher Scientific) in standard cell culture conditions (CO_2_ 5%, temperature 37°C with saturating humidity). Dulbecco’s Modified Eagle Medium (DMEM) with 10% of FBS (Fetal Bovine Serum) and 1% of penicillin, and streptomycin was used for cell culture medium. AlexaFluor568-pFN or AlexaFluor680-pFN (2 μg/ml) was added to the media on day 7 for 24 h after which cells were washed and fixed with 3.7% formalin for 10 min at RT followed by permeabilization with 0.25% Triton X 100 for 10 min. Cells were then blocked with 2% BSA. DAPI was used to stain the nuclei. Fluorescence images captured with AxioImager M2 microscope and Orca Flash 4.0 camera (Zeiss).

### pFN labeling and injections and detection in tissues

pFN (FC010, Sigma) was labelled with AlexaFluor 680 (AF680) as described above. Two groups of 12- week-old C57/bl6 male mice (n=3 for each group) (The Jackson Laboratory, Bar Harbor, ME, USA) were used. The experimental group received 20 ml/kg of AF680-pFN solution (i.e., 1mg pFN) via intraperitoneal (IP) injections while the control group was injected with inactivated AF680 that corresponded to the degree of pFN labelling (3.5 AF680 per pFN). Inactivation was done by mixing the AF680 with 0.5 M NaCl in 50 mM Tris, pH 8. This injection regimen was repeated on three consecutive days, with alternating injection sites. After 24 hours following the final injection, the mice were euthanized using CO_2_ asphyxiation under isoflurane anesthesia and following tissues were collected: plasma, liver, VAT (epididymal fat =visceral, VAT); SAT (inquinal fat = subcutaneous, SAT); BAT (brown fat), pancreas, BM (bone marrow) and SKM (quadricep femoris, skeletal muscle). SAT and VAT were imaged immediately *ex vivo*. All tissues were stored at –80 C for further analyses. *Ex vivo* epifluorescence imaging of was done with Perkin Elmer IVIS Spectrum In Vivo Imaging System (Perkin-Elmer, Waltham, MA, USA) at Mouse Housing Facility of The Rosalind and Morris Goodman Cancer Institute at McGill University. For fluorometric quantification, tissues were extracted with sodium dodecyl sulphate (SDS) and deoxycholic acid (DOC) and TritonX-100 containing buffer for maximal FN solubilization (39, 40)(150mM NaCl, 10mM Tris-HCl, 5mM EDTA, 0.1 % SDS, 1.0 % Triton X-100, 1.0 % Na-Deoxycholate PH 7.2). The fluorescence intensity in extracts was measured with Tecan Fluorometric Plate Reader Infinite 200 (Tecan, Männedorf, Switzerland) with Ex/Em 560nm/610 nm and expressed as mg of AF680-pFN per mg of total extracted protein (measured by BCA assay). The quantity of AF680-pFN was determined via the AF680 standard curve considering a 3.5/1 labeling degree. Plasma AF680-pFN gels were imaged with Molecular Dynamics / Amersham Typhoon 8600 Variable Mode Imager (Molecular Dynamics / Amersham Biosciences, Amersham, UK) or Odyssey M Imaging System (Li-Cor, Lincoln, NE, USA).

### Diet-induced obesity model

Diet-induced obesity model was generated from male and female Fn1-/-ALB mice and their litter-mate, sex-matched control *Fn1*flx/flx mice (n=5-11) with a 20 week special feeding regime of *ad libitum* HFD (41) (Envigo: 60% fat/lard and soybean oil), or nutrition-matched control diet (CD) (Envigo, 10% fat). Weight of the mice were assessed weekly. At the end point mice were assessed for insulin and glucose tolerance and fat mass (as below) and subcutaneous adipose tissue (SAT)(inquinal fat) and visceral adipose tissue (VAT) (epidydimal or periovarian) were collected for further analysis.

### Computed Tomography (CT) scans and Dual-energy X-ray absorptiometry (DEXA)-analysis for in vivo fat mass

All CT scan experiments were performed on nanoScan SPECT/CT for small animals (Mediso Medical Imaging Systems, Budapest, Hungary) at Small Animal The Small Animal Imaging Labs (*SAIL*) at Research Institute of McGill University Health Centre. The CT scans were conducted under anesthesia (Isoflurane 2%, medical air (0.6 L/min) delivered by a nose cone. Temperature and heart rate were monitored throughout the procedure using a Mediso system. The protocol was performed using 480 number of projections, scan method helical, 50 kVp, 610 µA, expose time 8 minutes and voxel size 250 µm. The reconstructed CT image was segmented automatically into the visceral adipose tissue (VAT) and the subcutaneous adipose tissue (SAT) using a MATLAB based software developed in house. MATLAB’s Canny algorithm was used to enhance the edge of the wall in order to separate the VAT and SAT into two independent volumes. In the case this failed for some animals, when the image of the abdominal muscular wall was not completely recognisable by MATLAB, we used the software *fsleyes* to conduct the segmentation along the muscular wall manually. Fat mass (%) was also analyzed with Dual-energy X-ray absorptiometry (DEXA) analysis at the Centre for Bone and Periodontal Research at McGill University. The DEXA scans were conducted under ketamine anesthesia using Lunar PIXImus II Mouse Densitometer (GE Medical Systems, Madison, WI) with 0.3mm stationary anode x-ray tube and following parameters: Filtration Nom: 2.5 mm Al(@70kV), Focal spot size: 0.25 mm x 0.25mm; Imaging area 100mm x 80mm; Max. X-ray tube voltage: 80kV; Max. X-ray tube current: 500 μA. After scanning, the fat% was analyzed by using custom-defined ROI function.

### Glucose tolerance test (GTT) and insulin tolerance test (ITT)

GTT and ITT were done at the endpoint of the experiment at 20 weeks on CD or HFD groups after 6 hours of fasting. For the GTT, mice were given IP injections of glucose (1 g/Kg body weight) and blood glucose level was measured using a glucometer (FreeStlye Life, Abbot) at 0, 15, 30, 60, and 120 min after glucose injection. For the ITT, insulin (0.8 U/Kg body weight) (Humulin® N Lilly. Canada) was injected, and blood glucose levels were measured at 0, 30, 60, 90, and 120 min after insulin injection. The area under the curve was calculated using GraphPad Prism vs 10.

### In vivo and in vitro insulin sensitivity of adipose tissue

Insulin sensitivity of SAT and VAT was conducted via *in vivo* insulin injections and quantification of phosphorylation ratio in signalling pathways via Western blotting. *Fn1*flx/flx and *Fn1*-/-ALB mice on a CD and HFD were fasted for 6 h and given IP injections of insulin (1-2 U/kg) (Humulin® N Lilly. Canada) Mice were euthanized 15 min post-injection followed by dissecting inguinal fat, i.e., subcutaneous adipose tissue (SAT) and perigonadal fat; epididymal (male) and periovarian (female) representing visceral adipose tissue (VAT) and homogenized in extraction buffer. The extraction buffer contains 100 mM Tris (pH 7.4), 1% Triton X-100, 10 mM EDTA, 100 mM sodium fluoride, 2 mM phenylmethyl sulfonyl fluoride, 5 mM sodium orthovanadate, in addition to phosphatase inhibitor and protease inhibitor cocktail. The protein lysate was analyzed by Western blotting as described below. The blots were probed with anti-Phospho-AKT (Ser473 and Thr308)/anti-total-AKT. Quantification of band intensities was done with the NIH Image J open-source image processing program.

### Quantitative RT-PCR (qPCR) for adipocyte markers

mRNA was extracted using the RNeasy Mini Kit (Qiagen, Venlo, Netherlands) and cDNA prepared using the High-Capacity cDNA Reverse Transcription Kit (Applied Biosystems, Foster City, CA). qPCR was done on the StepOnePlus Real-Time PCR System (Applied Biosystems) in a 20 µl reaction volume consisting of the following: 9 µl (50 ng) synthesized cDNA, 10 µl TaqMan Fast Advanced Master Mix and 1 μl of each TaqMan Gene Expression Assay. TaqMan® Fast Advanced Master Mix and primers were purchased from Applied Biosystems. Following primers were used: *Pref1* (Mm00494477_m1), *Pparg* (mM00440940_m1), *Cebp* (Mm00514283_s1), *Adipoq* (Mm00456425_m1), *Ucp1* (Mm01244861_m1), *Prdm16* (Mm00712556_m1). Expression was normalized to Gapdh (Mm99999915_g1).

### Leptin and adiponectin

Plasma was collected at end point of each feeding regime after 6 h fasting. Leptin and adiponectin levels were measured with Mouse Leptin ELISA Kit (ab100718) and Mouse Adiponectin ELISA Kit (ab108785) (Abcam, Cambridge, UK) according to the instruction of the manufacturer.

### Protein extraction and Western blotting

Protein extractions were prepared using tissues were extracted with sodium dodecyl sulphate (SDS) and deoxycholic acid (DOC) and TritonX-100 containing buffer for maximal FN solubilization (39, 40)(150mM NaCl, 10mM Tris-HCl, 5mM EDTA, 0.1 % SDS, 1.0 % Triton X-100, 1.0 % Na-Deoxycholate PH 7.2) containing a protease inhibitor and phosphatase inhibitor cocktails (Sigma). Protein concentrations were determined using the BCA protein assay kit (Pierce, Rockford, IL, USA). For Western blotting, proteins were resolved by 8.5% SDS-PAGE gel electrophoresis and transferred to PVDF membranes (Bio-Rad, Mississauga, ON, Canada). Membranes were blocked with 5% non-fat milk powder in Tris-buffered saline-Tween (TBS-T) buffer, and each protein was detected with specific antibodies followed by corresponding HRP-conjugated secondary antibodies. The detection was visualized with Amersham ECL Prime Western Blotting Detection Reagent (GE Healthcare Life Sciences, Baie d’Urfe, QC, Canada). Quantification of band intensities (where indicated) was done with the NIH Image J open-source image processing program.

### Adipose tissue histology

At the endpoint of HFD and CD feeding *Fn1*flx/flx and *Fn1*-/-ALB mice SAT and VAT were collected for histological analysis. Tissues were fixed in 3.7% formalin, paraffin-embedded, sectioned and stained with hematoxylin & eosin (H&E) staining. The number and size of cells were quantified using open-source image analysis software, CellProfiler (https://cellprofiler.org/) (42, 43) via a systematic assessment of cell size distribution from 5 separate fields of microscopic image per genotype. The workflow using CellProfiler or ImageJ, with a focus on the “*Adipocytes_Sizing_Pipeline.”* In the CellProfiler pipeline, the *ColorToGray* module initially converted the colour images to grayscale, followed by the transformation into a binary black-and-white representation using the *ImageMath* module. Subsequent modules, including *EnhancedOrSuppress* Features, *EnhanceEdges*, and Morph, were strategically applied to enhance membrane visibility and facilitate adipocyte distinction by eliminating specks and sharpening edges. The *IdentifyPrimaryObjects* module was used to delineating adipocyte membranes based on the enhancements achieved in prior steps. The subsequent *CovertObjectsToImage* module integrated these identified objects back into the original black-and-white image, resulting in a clearer representation of the cell membranes. To specifically focus on adipocytes, another round of processing was conducted using *ImageMath* and EnhancedOrSuppressFeatures modules. The *IdentifyPrimaryObjects* module was then reapplied, this time recognizing adipocytes as objects. In *Test Mode*, the module allowed visual confirmation of adipocyte identification, providing flexibility for manual adjustments to meet specific preferences.The *MeasureObjectSizeShape* module quantified key parameters such as the area of each adipocyte, generating raw measurements in pixels. To convert these measurements into microns, the *CalculateMath* module was utilized, incorporating a scaling diagram approach. This involved determining the pixel-to-micron conversion factor using the scaling diagram present in each image. Aa 10X magnification, the conversion factor was calculated as 0.13516 μm^2^/pixel^2^, enabling accurate transformation of pixel-based area measurements into meaningful micrometre values. Adipose cell size was reported based on the surface area of detected cells. According to previous studies the adipocyte exhibits a comparable spherical surface area within the range of 5,000 to 1,000,000 μm² (44).

### RNA sequencing and data analyses

The total RNA was extracted from SAT of CD-fed pFN KO mice and their control group of flx/flx on CD and HFD-fed VAT tissue of flx/flx and pFN KO. Total RNA was quantified, and its integrity was assessed on a LabChip GXII (PerkinElmer) instrument. Libraries were generated from 250 ng of total RNA as following: mRNA enrichment was performed using the NEBNext Poly(A) Magnetic Isolation Module (New England BioLabs). cDNA synthesis was achieved with the NEBNext RNA First Strand Synthesis and NEBNext Ultra Directional RNA Second Strand Synthesis Modules (New England BioLabs). The remaining steps of library preparation were done using and the NEBNext Ultra II DNA Library Prep Kit for Illumina (New England BioLabs). Adapters and PCR primers were purchased from New England BioLabs. Libraries were quantified using the KAPA Library Quanitification Kits - Complete kit (Universal) (Kapa Biosystems). Average size fragment was determined using a LabChip GXII (PerkinElmer) instrument. The libraries were normalized and pooled and then denatured in 0.05N NaOH and neutralized using HT1 buffer. The pool was loaded at 175pM on a Illumina NovaSeq S4 lane using Xp protocol as per the manufacturer’s recommendations. The run was performed for 2×100 cycles (paired-end mode). A phiX library was used as a control and mixed with libraries at 1% level. Base calling was performed with RTA v3.4.4. Program bcl2fastq2 v2.20 was then used to demultiplex samples and generate fastq reads. The data was analyzed with R Statistical Software (https://www.r-project.org/)(45) in R Studio and results were plotted using ggplot2, ggrepel and dplyr libraries from comprehensive R archive network (CRAN). Differentially expressed genes (DEGs) were analyzed between pFN KO and flx/flx mice under a normal diet (SAT). Transcripts were considered differentially expressed when the absolute value of the log-fold change was greater than 2, and the p-value was less than 0.05. Gene enrichment analysis was done with Metascape (https://metascape.org) and selected overrepresented GOterms were confirmed via analysis using Panter database (https://www.pantherdb.org/) *and* KEGG (Kyoto Encyclopedia of Genes and Genomes (https://www.genome.jp/kegg/).

### Statistical and data analysis

Data were analyzed with GraphPad Prism software (version 10). Results were expressed as mean ± SEM (standard error of the mean). Data were analyzed by the one-way and two-way analysis of variance (ANOVA) followed by multiple comparison Tukey post-hoc test. When comparing two groups Student’s Ttest was used. The differences were considered statistically significant for p-values < 0.05. (*p < 0.05, **p < 0.01, ***p < 0.001, ****p < 0.0001, ns: not significant).

## 3. Results

pFN circulates in blood at high concentration and is known to contribute to ECM of several tissues. In this study we have examined its potential contribution to AT and metabolism. We first examined if circulating pFN accumulates from blood to metabolic tissues; VAT, SAT, BAT, liver, pancreas, bone marrow and skeletal muscle via labeling pFN with AlexFluor568 (AF568-pFN) and AlexaFluro680-pFN (AF680-pFN) followed by intraperitoneal (IP) injections to normal, adult mice. Prior to IP injections we confirmed in 3T3- L1 preadipocyte cell cultures that the fluorescence labelling does not interfere with pFN ability to create fibrillar matrix and that the inactivated AF-label alone does not integrate to cell layers (**Figure 1A**). The *in vivo* study involved an injection regime of three IP injections in consecutive days as we and others have done before (**Figure 1B**)(27, 28, 46). Control injection group included inactivated AF568 or AF680 label that corresponded to the labeling degree calculated for AF-pFN. After the three injections, intact full length AF568-pFN was detected in plasma by fluoroimaging of plasma run into SDS-PAGE gels (**Figure 1C**). At the end of the experiment, mice were euthanized, plasma was collected and metabolic tissues (plasma, liver, VAT (epididymal fat =visceral, VAT); SAT (inquinal fat = subcutaneous, SAT); BAT (brown fat), pancreas, BM (bone marrow) and SKM (femoral quadriceps) were dissected, and proteins extracted using SDS-DOC-TritonX100 containing buffer to solubilize ECM. AF568 fluorescence was measured with fluorometer and expressed as μg of pure labelled pFN-568 per mg of tissue (determined from a standard curve) (**Figure 1D**). Significant 568 fluorescence in plasma (compared to control inactivated AF568 injected groups) was detected and high levels of AF568-pFN accumulation to liver, SAT, VAT and bone marrow (BM) was observed. BAT, pancreas and skeletal muscle (SKM) also showed significant AF568-pFN accumulation but markedly less than others assessed (**Figure 1D**). AF568-pFN accumulation in VAT and SAT were also confirmed with *ex vivo* epifluorescence imaging (**Figure 1E**) and via SDS-gels of the extracts followed by fluoroimaging (**Figure 1F**). These data demonstrate that pFN is a component of AT depots and suggests a role in AT function.

**Figure.1.**
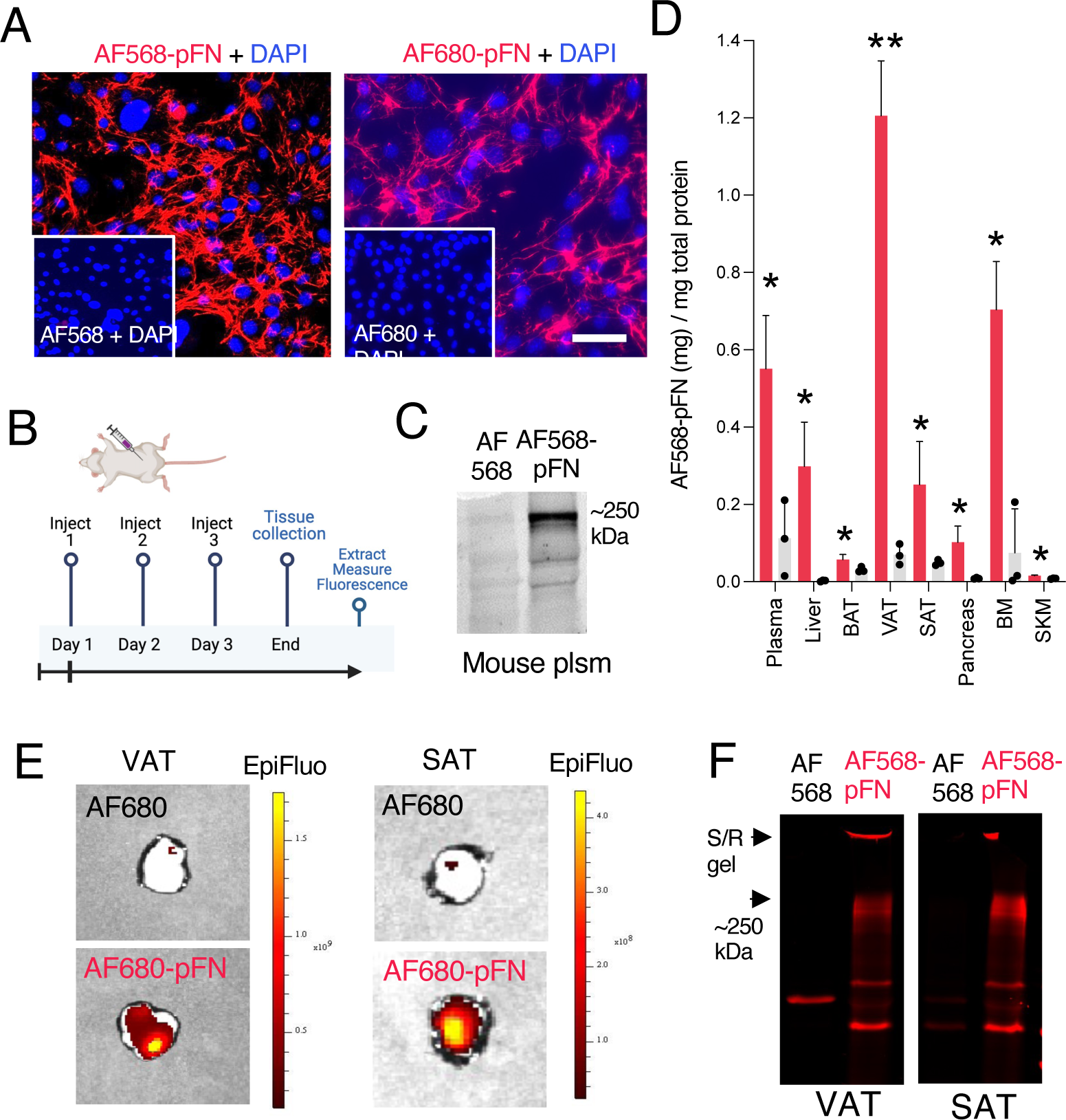
pFN accumulates from circulation to metabolic tissues. **A.** pFN was labelled with AlexaFluor568 (AF568) and with AlexaFluor680 (AF680) to confirmed to undergo fibrillogenesis in MEF cell cultures. Inactivated AF-fluorophores were used as controls and did not show labeling in the cultures. Magnification bar equals 20 μm. **B.** Injection regime involved three consecutive IP injections of 1 mg of AF568-pFN per day. Control mice received inactivated AF568 corresponding to the labeling degree of AF568-pFN (3.5 AF568 in each pFN). Created with BioRender.com. **C.** Mouse plasma (20 mg total protein) analyzed by SDS-PAGE and visualized by fluoroimager shows presence of AF568-pFN circulation after injections. **D.** Plasma, liver, VAT (epididymal fat =visceral, VAT); SAT (inquinal fat = subcutaneous, SAT); BAT (brown fat), pancreas, BM (bone marrow) and SKM (quadricep femoris, skeletal muscle) levels of AF568-pFN were quantified after SDS-DOC-TritonX100 extractions of the tissues followed by fluorometric analyses. Graph demonstrates accrual of pFN to all metabolic tissues, highest levels detected in VAT. Plasma (plsm) visualized as control and output. Error bars represent SEM. n = 3 (WT male mice). Statistical significance represents AF568-pFN vs AF568. *p < 0.05, **p < 0.001. **E.** *Ex vivo* imaging AF680-pFN accumulation to VAT and SAT. Imaging was done with IVIS Spektrum immediately after dissecting the tissues. **F.** Accumulation of AF568-pFN was further visualized via SDS-PAGE gels of SAT and VAT extracts. Full length AF568-pFN is detected at 250 kDa with some high molecular weight material at the border of stacking and running gel (S/R gel). Error bars represent SEM (n=3 mice per group). Statistical significance is represented as: *p < 0.05.

To investigate of pFN affects adipocytes or AT expansion, i.e., and weight gain and in whole-body metabolism, we developed diet-induced obesity mouse model in the absence of plasma FN. For this, pFN deficient mice were created via breeding *Fn1*flx/flx and ALB-Cre (47) (**Figure 2A**). These mice have been shown to have a 96% decreased of the normal circulating plasma fibronectin levels (47, 48). Our model showed clear decrease in FN in plasma particularly in male mice, and otherwise similar protein pattern in SDS-PAGE analysis (**Figure 2B**). pFN KO mice and their littermate controls (flx/flx)(in Figures referred to as KO and flx, respectively) of both sexes, were placed on high-fat diet (HFD) and nutrition-matched control diet (CD) for 20 weeks with weekly weight monitoring. At the end point, FN levels in plasma and adipose tissues (VAT and SAT) were analyzed for FN (general antibody that detects all FN, including cellular FN) showing clear absence of circulating FN in pFN KO on both diets. The pFN KO had similar circulating FXIII- A levels (**Figure 2C**). Interestingly, the pFN KO SAT was essentially all depleted in the pFN KO demonstrating that majority of AT FN is from circulation (**Figure 2C)**. Furthermore, FXIII-A transglutaminase levels showed also a clear decrease suggesting a regulatory a link between two. Total FN levels in VAT were not as clearly affected as in SAT for neither FN or FXIII-A.

**Figure 2.**
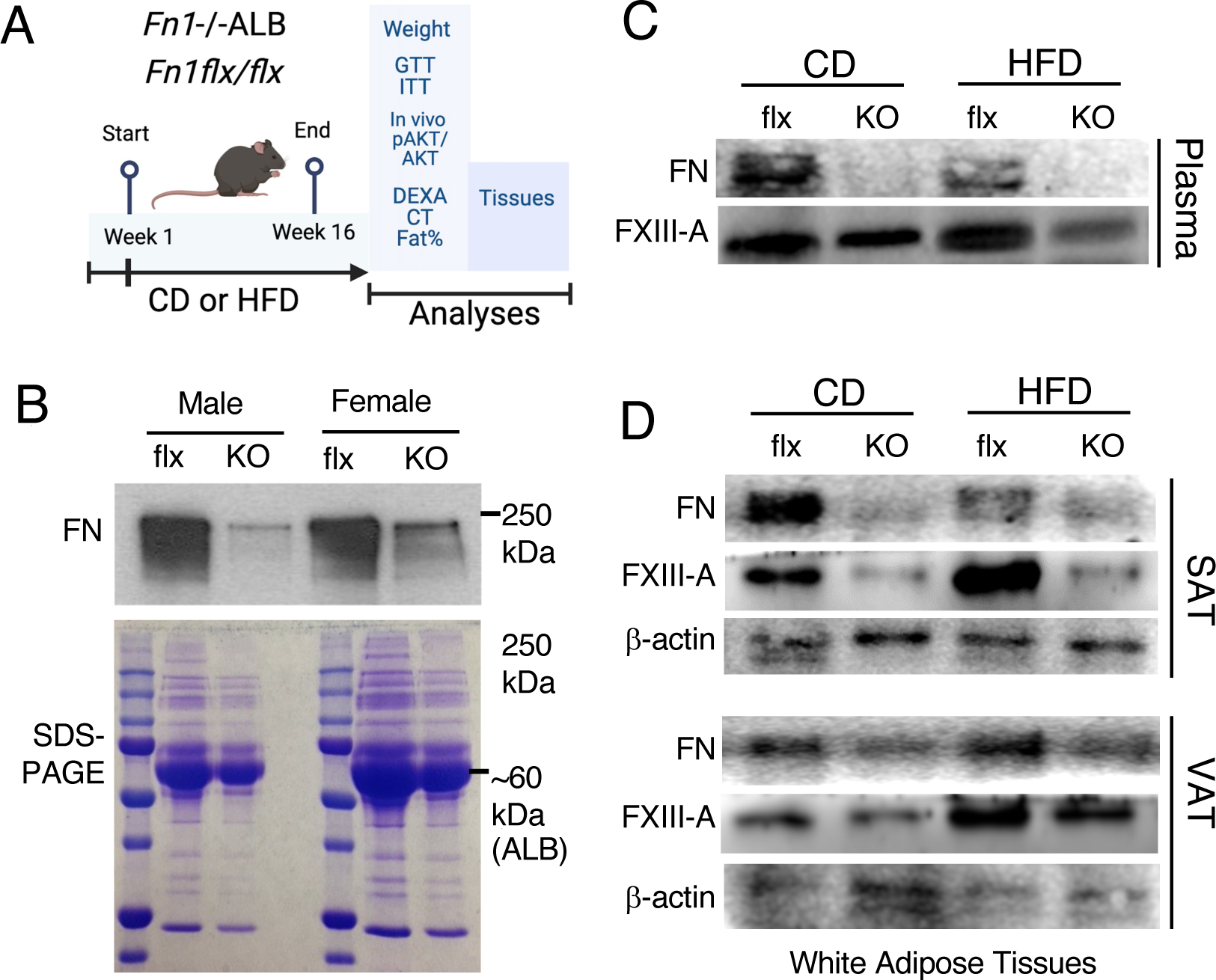
Liver-specific (ALB-Cre) *Fn1* knockout mice, aka, plasma FN knockout mice show no or low FN levels in white adipose tissues. **A**. *Fn1*-/-ALB; pFN knockout mice (referred to as KO) were developed by breeding of *Fn1*flx/flx and ALB-Cre mice. Mice were fed obesogenic high-fat diet (HFD) for 20 weeks which was followed by metabolic and adipose tissue phenotyping. Control mice were fed nutrition-matched control diet. Created with BioRender.com. **B.** Analysis of plasma shows efficient elimination of circulating FN, particularly in male mice compared to control (flx/flx)(referred to as flx). Females appear to have residual circulating FN. Lower panel shows Coomassie Blue stained SDS-PAGE of whole plasma protein panel (0.5 mg total protein loaded) where albumin (ALB) shows a strong band. **C** pFN and FXIII-A levels in plasma and WAT (inguinal WAT; Subcutaneous AT, SAT) (epididymal WAT; Visceral AT; VAT) in normal, control diet and HFD-fed mice. Neatly all SAT FN is eliminated with simultaneous low FXIII-A levels in the tissue. VAT FN and FXIII-A levels are less affected.

Weight and fat mass analysis at the end point of the 20-week feeding demonstrated that pFN KO mice did not have altered weight gain compared to their control mice on HFD (**Figure 3A**). Female mice, pFN KO or flx/flx, did not show significant weight gain on HFD and thus further steps of the study was conducted mostly with male mice. Fat mass analysis of male mice was measured via dual X-ray absorptiometry (DEXA) scans, pFN KO on CD had notably lower total fat mass % albeit not significant (p=0.058) (**Figure 3B**). Whole-body CT (computed tomography) scans SAT and VAT depots showed clear effect of HFD on both depots but no statistical difference between pFN KO and controls (SAT HFD: p= 0.1249 and VAT HFD; p=0.5877) (**Figure 3C,D**). Reflectin the weigh and AT mass, circulating leptin and adiponectin levels were not altered between flx and pFN KO (**Supplemental Figure 1**).

**Figure 3.**
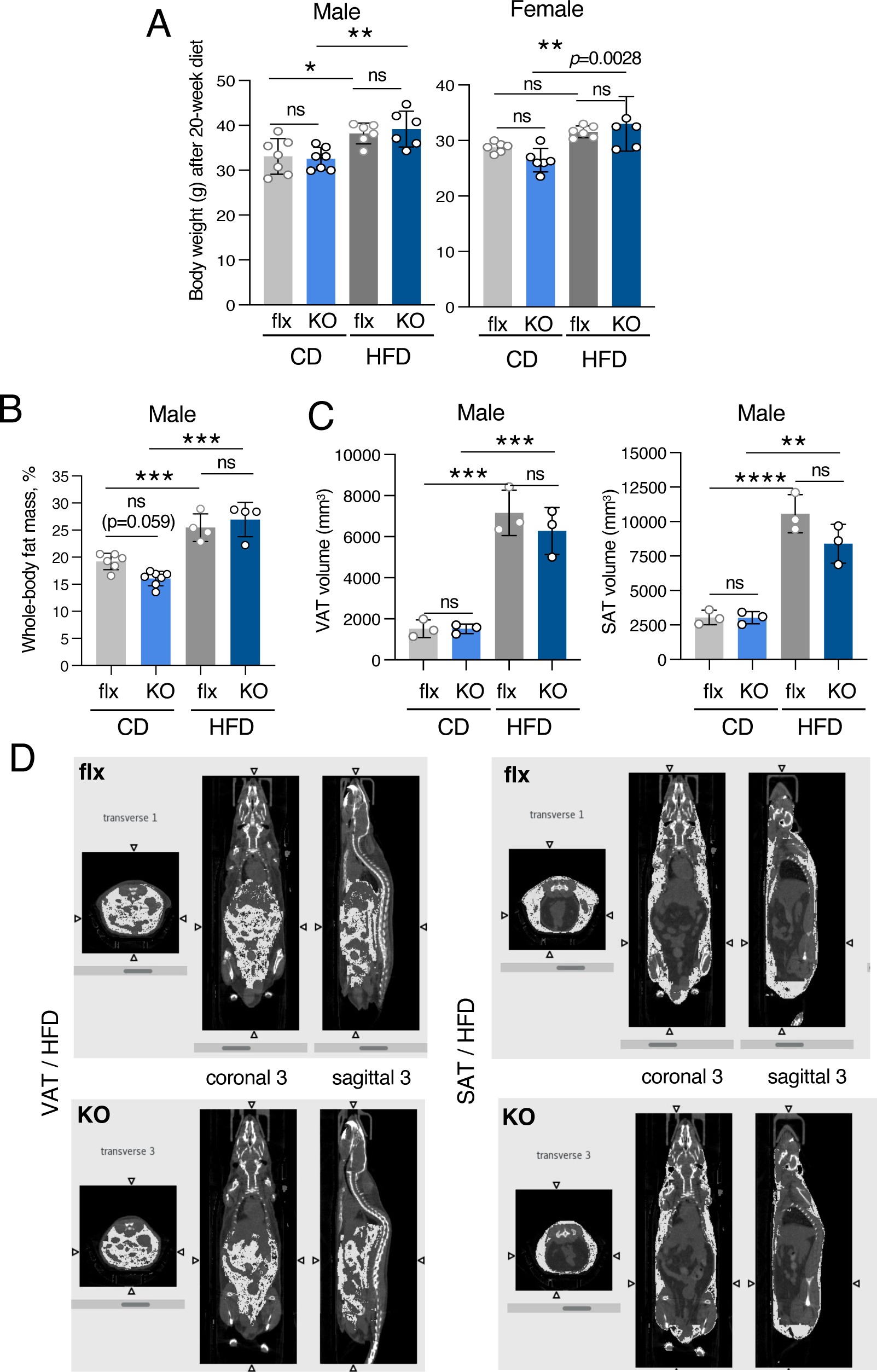
pFN knockout mice gain normal weight on high fat diet (HFD) and have normal levels of whole-body fat, VAT and SAT. **A.** Body weights of male and female pFN KO and its control flx mice after 20 weeks on control diet (CD) and high fat diet (HFD). Female mice did not show significant weight gain after HFD feeding in control mice. **B.** Dual-energy X-ray absorptiometry (DEXA)-scans show no significant difference between male flx and KO mice in whole body fat mass (%). **C.** Computed tomography (CT) scans for visceral (VAT) and subcutaneous (SAT) adipose tissue show no significant difference between male flx and KO after 16-week HFD. CT scan images (transverse, coronal and sagittal images show equal and similar fat distribution in KO and flx mice after HFD. Error bars represent SEM (n=3-5 per group). Statistical significance is represented as: *p < 0.05, **p < 0.001, ***p < 0.0001, ****p < 0.00001.

Analysis of fasting glucose at the end point of the 20-week feeding showed no significant differences in male or female, flx/flx versus pFN KO mice (**Supplemental Figure 2**). However, glucose tolerance test (GTT), surprisingly, showed significantly better glucose clearance in male pFN KO mice compared to flx mice on *normal CD-feeding* (**Figure 4A**). This difference disappeared on HFD (**Figure 4A**). Female mice did not show differences (**Supplemental Figure 3**). Similarly, insulin tolerance test (ITT) to probe whole-body insulin sensitivity showed significant enhancement in male pFN KO mice compared to flx/flx mice *on normal CD-feeding* (**Figure 4B**). Also, this difference disappeared on HFD. No changes were observed in female mice (**Supplemental Figure 4**).

**Figure 4.**
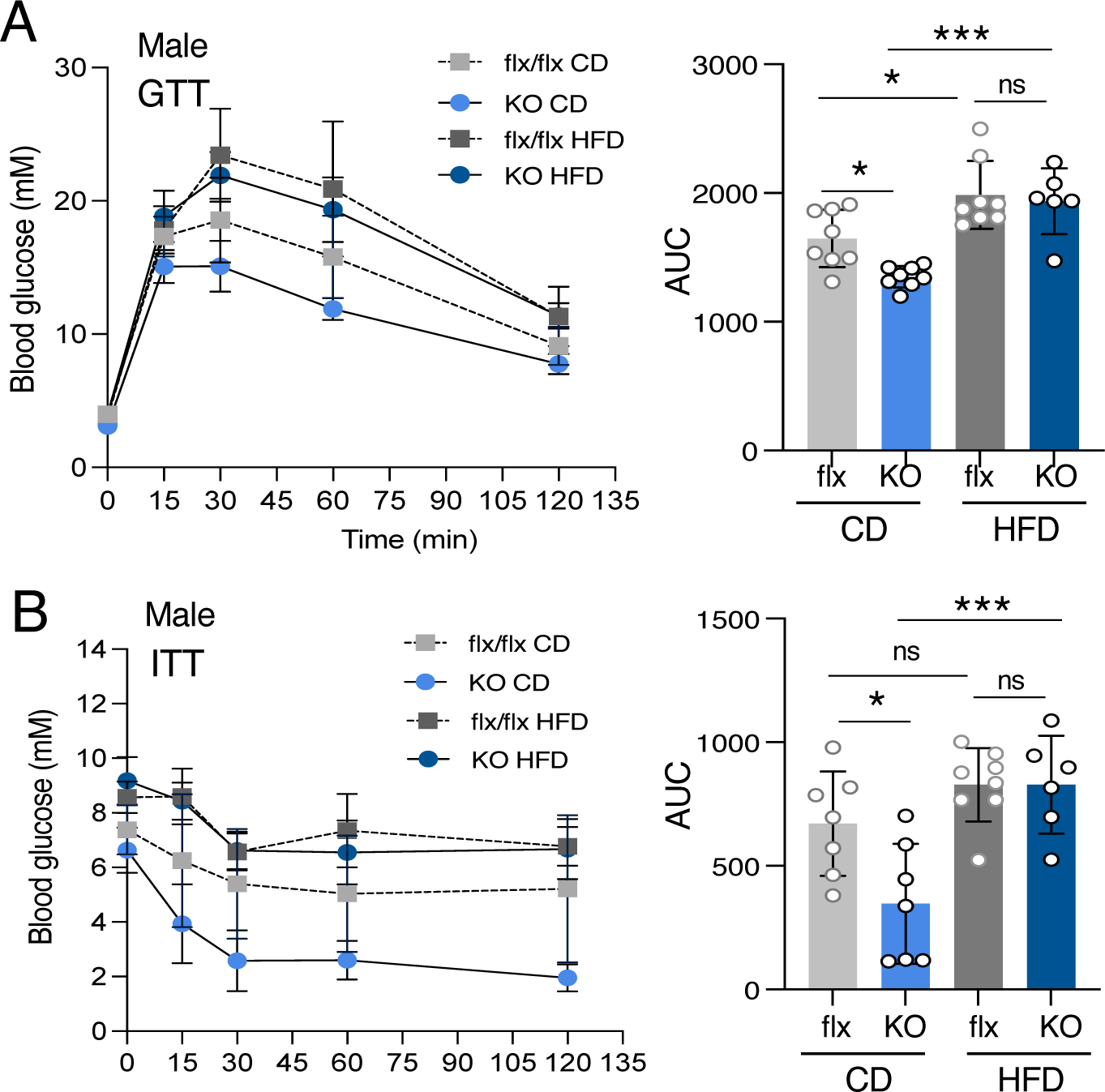
Whole-body glucose clearance and insulin sensitivity is enhanced in male pFN knockout on normal diet. **A.** At the end point of the 20-week CD and HFD-feeling, pFN KO (KO) and control flx/flx mice were tested for glucose clearance via classic glucose tolerance test (GTT) Significantly enhanced glucose clearance was detected in pFN KO mice on CD which presented significantly smaller AUC values. The difference disappeared in HDF. **B.** Insulin tolerance test (ITT) was conducted via injections of bolus of insulin (ITT) followed by glucose clearance expressed by Area Under the Curve (AUC). pFN KO on CD mice showed significantly smaller AUC, i.e., enhanced insulin sensitivity. Error bars represent SEM (n=6-8 per group). Statistical significance is represented as: *p < 0.05, **p < 0.001, ***p < 0.0001.

Interestingly, *in vivo* insulin sensitivity assay of SAT and VAT with bolus of insulin injections followed by rapid dissections, tissue extraction and signaling analysis by WB, supported the observed differences on normal diet. pFN KO SAT after *normal CD-feeding* showed significantly increased phosphorylation of AKT to both Ser473 and Thr308 (**Figure 5A)** which disappeared on HFD (**Figure 5**). VAT insulin sensitivity, pAKT(Ser473)/AKT was similarly significantly increased in the pFN KO and the effect remained significant also on HFD (**Figure 5B**). No phosphorylation to Thr308 was detected in VAT.

**Figure 5.**
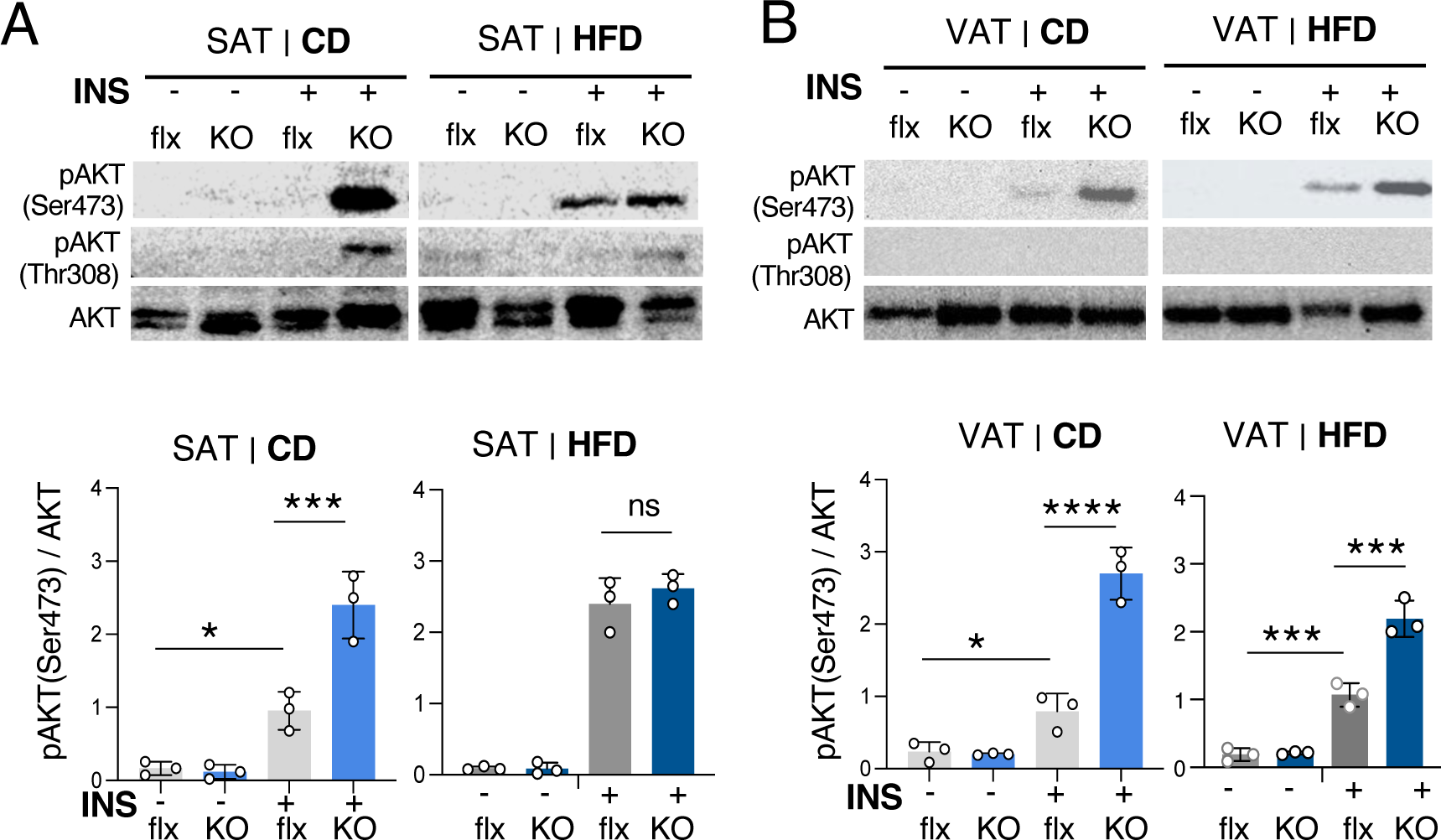
Absence of pFN significantly increases *in vivo* insulin sensitivity of SAT and VAT on CD normal SAT. pFN KO and flx/flx mice, after a 20-week CD and HFD-feeding, were assessed for *in vivo tissue* insulin sensitivity at the end point via injections of a bolus of insulin (INS)(+), which was followed by rapid tissue dissections and extraction 15 min later. Signaling induction was was assessed via Western blotting for pAKt(Ser473 and Thr308). Control groups were injected with saline (-). WBs were quantified by Image J image analysis. **A.** pAKT (Ser473 and Thr308) were both engaged in SAT upon insulin injections on CD tissue. This difference was significant and disappeared on HFD. **B.** Significantly increased pAkt/(Ser473)AKT was also detected in VAT, but no Thr308 phosphorylation was observed in CD or HFD groups. Interestingly, increased insulin sensitivity of VAT remained significantly increased also in the HFD group. Error bars represent SEM. n = 3. Statistical significance is represented as: *p < 0.05, **p < 0.001, ***p < 0.001, ****p < 0.0001.

To get insight also at tissue level how absence of pFN affects AT and insulin sensitivity, histological analysis of SAT and VAT was done. Analysis revealed visibly smaller cells in pFN KO mice on *normal feeding (CD)* which was confirmed significant by cell size quantification. Both normal CD-fed pFN KO SAT an VAT had significantly increased numbers of small adipocytes (**Figure 6A and B**) with SAT specifically showing pockets of high cellularity (**Figure 5A, inset**). HFD-feeding eliminated the small adipocytes and showed in fact significantly less small adipocytes than flx/flx controls suggesting some alterations adipogenesis and/or hypertrophy.

**Figure 6.**
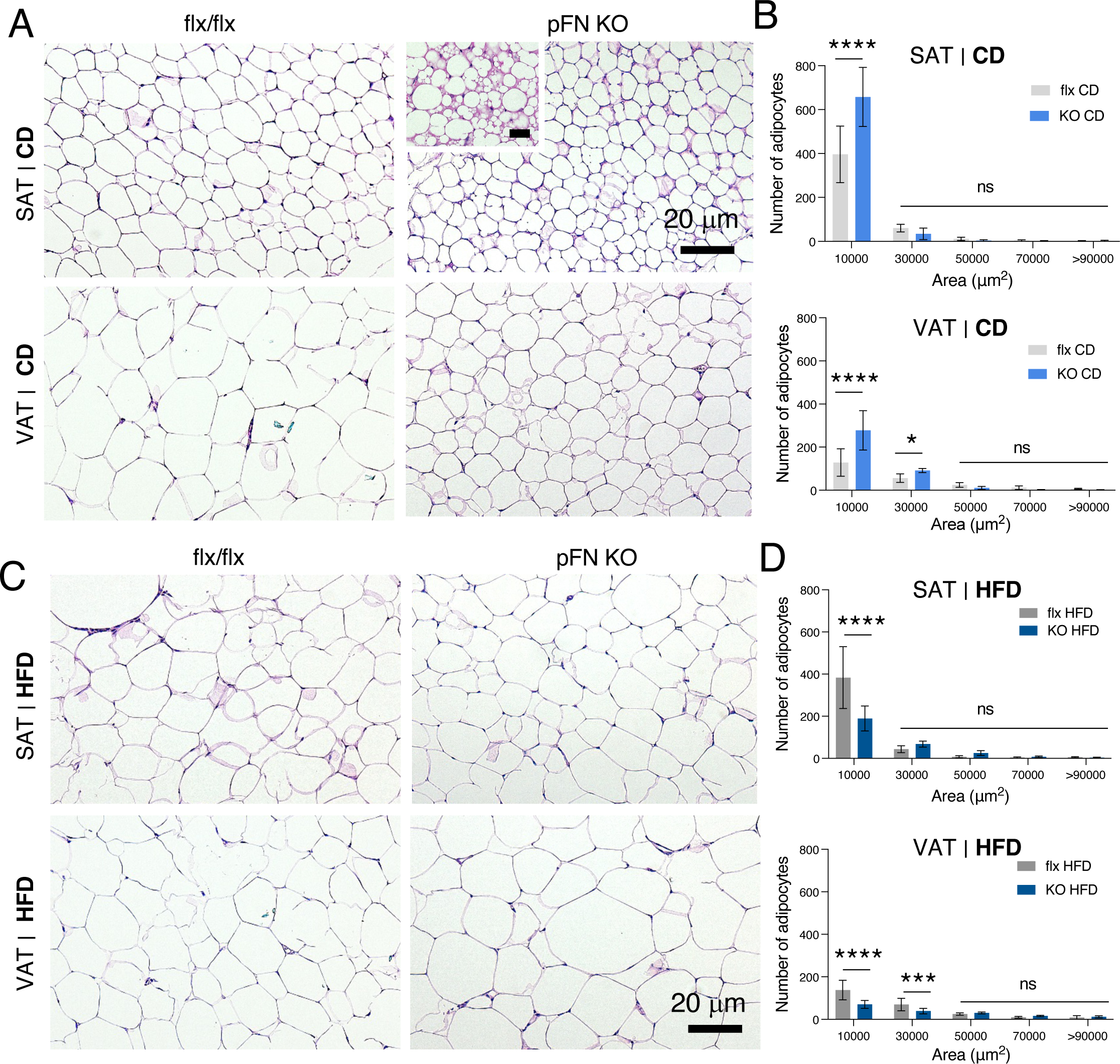
Histological analysis SAT And VAT shows increase in smaller adipocytes in pFN KO on control diet. **A.** H&E stained histological sections of SAR and VAT on CD shows visible smaller adipocytes in pFN KO. In SAT, small pockets of high cellularity wer observed (insert). Magnification bar; 20 μm **B.** Adipocyte size was quantification using CellProfiler program and grouping cell numbers per size/ adipocyte area, μm^2^. confirms the significant increase in smaller adipocytes. **C.D** HFD-feeding visibly decreased the differences in cell size as seen by H&E stained histological sections. Cell size assessment (adipocyte area, μm^2^) shows significantly decreased amount of smaller cells in both SAT and VAT in pFN KO compared to flx/flx control. Error bars represent SEM (n=5 mice per group). Statistical significance is represented as: *p < 0.05, ***p < 0.0001, ****p < 0.00001.

pFN KO effect on SAT phenotype on CD and VAT phenotype on HFD diet were most profound and we explored possible mechanisms for increased insulin sensitivity and cellularity further via RNA sequencing. SAT had a total of 85 Differentially Regulated Genes (DEGs) with a cut-off criteria of 2-fold change with False Detection Rate (FDR)-significance of *p*=0.05 (**Figure 7A**). Gene enrichment analysis using Metascape, and confirmation with Panther and KEGG, revealed three distinct and prominent Biological Processes: one on immunity marked by the abundant presence of immunoglobulins in SAT, and second on long-chain fatty acid metabolism and fatty acid biosynthesis, and third on positive regulation of cold-induced thermogenesis suggesting beiging of SAT (**Figure 7B**). Genes contributing to this Gene Ontology term included *Elovl3, Ucp1, Elovl6, Alox12e* of which *Ucp1* (Uncoupling Protein 1) is a key thermogenesis protein and showed 7-fold upregulation in the RNAseq data. These data prompted us to examine adipocyte differentiation markers by qPCR (**Figure 7C**) which showed significant increase in preadipocyte marker, *Pref1*, and also mature adipocyte markers *Pparg* and *Adipoq. Cebp* was not changed. Most notably, *Ucp1* was confirmed significantly increased, as was beige adipocyte precursor marker *Pdrm16* (**Figure 7D**) suggesting clear alterations in adipocyte pools, differentiation and beiging of SAT. HFD-fed VAT in turn had more dramatic alterations; 444 Differentially Regulated Genes (DEGs) of which 77% (342 were downregulated) and 102 upregulated. Gene enrichment analysis revealed several prominent gene functional clusters most interesting ones related to skeletal and limb development marked by downregulation of Hox-family transcription factors (*Hoxa10, Hoxa11, Hoxa9, Hoxb9, Hoxc10, Hoxa13, Hoxb6,Hoxb7,Hoxb8*) and skeletal stem cell-osteogenesis and mineralization markers (such as *Grem1; Runx2, Lef1, Enpp1, Pitx1, Bglap3, and Pthr1* (parathyroid hormone 1 receptor). Also, tube morphogenesis and branching morphogenesis of an epithelial tube were represented suggesting alteration in angiogenesis.

**Figure 7.**
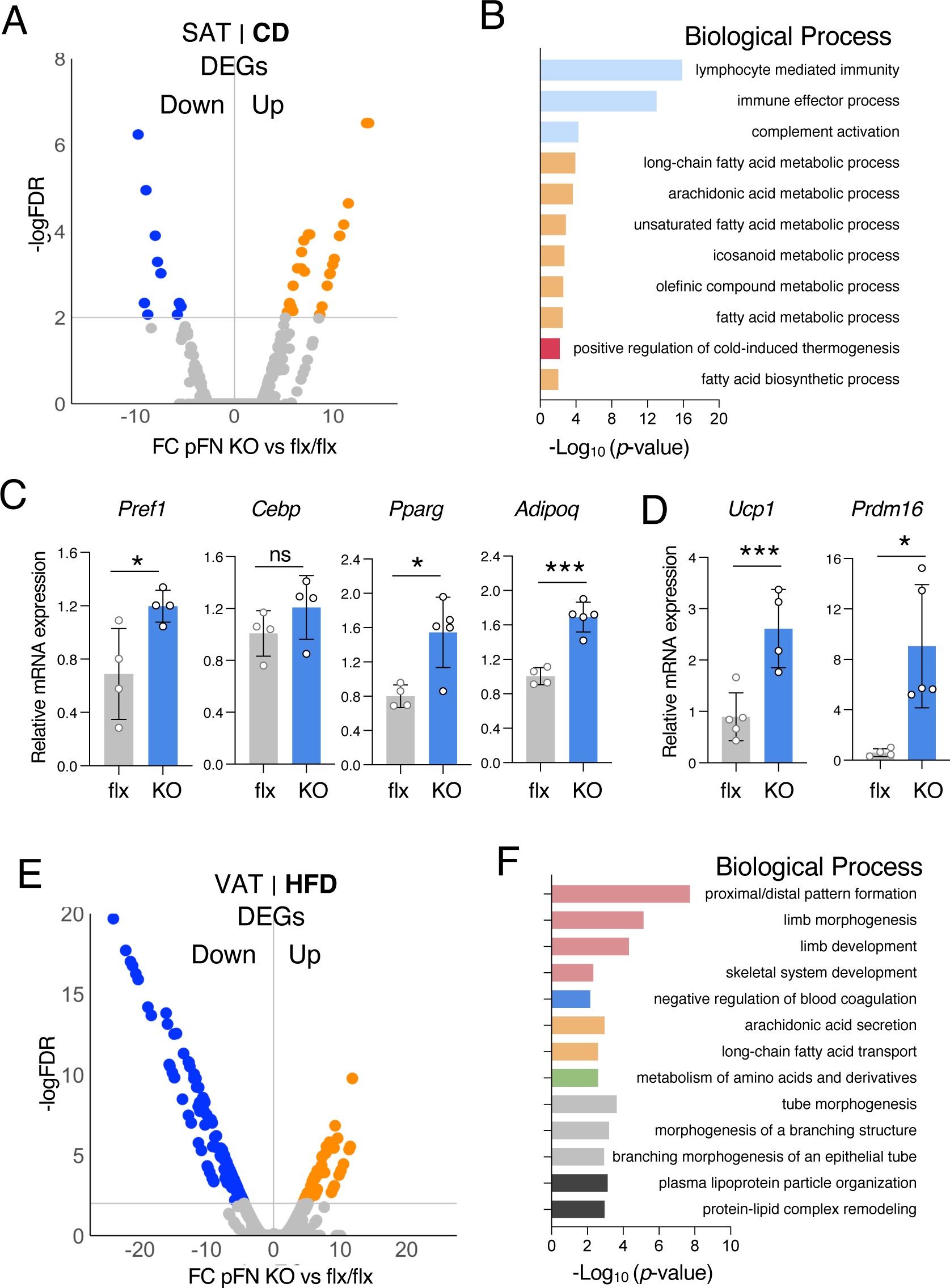
RNAseq and qPCR analyses of pFN vs flx SAT from normal, control diet and VAT from HFD feeding reveals significant chances in adipose tissue. **A.** RNA sequencing of SAT from CD-fed mice revealed 85 significant Differentially Regulated Genes (DEGs); 25 downregulated and 60 upregulated. Note that data points can contain more than one gene. **B.** Metascape analysis of gene enrichment of overrepresented genes revealed several pathways altered in normal pFN KO SAT, most notably relevant to adipocytes are changes in fatty-acid metabolic processes, and cold-induced thermogenesis. **C.** qPCR analysis of adipocyte markers in CD-fed pFN KO SAT showed significant increase in expression of *Pref* (preadipocyte) and *Pparg, Adipoq* (mature adipocytes), but not in (*Cebp*). **D.** Thermogenesis marker *Ucp1* showed significant increase along with *Prdm16* (beige adipocyte marker) in CD-fed pFN KO SAT. **E.** RNA sequencing of VAT from HFD-fed pFN KO and control mice revealed 444 significant Differentially Regulated Genes (DEGs); 342 downregulated and 102 upregulated. Note that data points can contain more than one gene. **F.** Metascape analysis of gene enrichment of overrepresented genes in HFD-fed pFN KO VAT revealed several altered pathways, most down regulated. The top ones were related to skeletal development marked by changed in transcription factors that guide commitment to osteogenesis. Error bars represent qPCR SEM. n = 4. Statistical significance is represented as: *p < 0.05, **p < 0.001, ***p < 0.001, ****p < 0.0001.

## 4. Discussion

ECM is an important regulator of adipogenesis and AT homeostasis with effects on whole body metabolism (49–53). Generally, ECM components are synthesized locally in AT by the adipocyte lineage cells as well as other cell types. Here we provide evidence that a circulating ECM molecule, pFN, accumulates to AT and contributes to the AT homeostasis and cell pools. The role of pFN as a major source of tissue ECM *in vitro* and *in vivo* was first introduced over 40 years ago by Oh et al. (25) and further elaborated in several studies including Sottile et al. (54) who demonstrated that *Fn1*-/- MEF cultures can assemble normal FN matrix from the serum pool of FN. A study by Moretti et al. (24) further demonstrated that mice with hepatocytes genetically engineered to produce EDA-domain in *Fn1* gene exhibited a significant reduction in FN levels in both plasma and tissues. Conversely, when liver-specific deletion of the EDA-domain was removed again in these same animals, FN levels in plasma and tissues (skin, kidney, liver, heart muscle, lung, and brain) were restored. This study emphasized pFN essential contribution to tissue ECM and showed that FN harboring EDA-domain cannot be secreted from hepatocytes (24). The development of *Fn1*flx/flx mouse (47) has allowed for further understanding of pFN in different systems via liver-specific *Fn1* knockout and its role has been linked to supporting cell survival and tissue integrity such as in brain injury (47), normal bone matrix (28), and during vascular development (27) but no roles in skin healing or hemostasis. Our data provides AT to this list as a tissue that accumulates pFN with important consequences to its function.

*In vitro* evidence has long suggested that total FN matrix (source was not defined, can be cFN or/and pFN) sustains preadipocyte phenotype and inhibits adipogenesis via binding to Pref1 (33). It was demonstrated that, FN fragments containing _I1-5_, such as 24k Fib I, promote adipogenesis by blocking FN fibrillogenesis and matrix assembly (29–31). Our data demonstrated that pFN from cell culture serum, jointly with transglutaminase FXIII-A from pre-adipocytes, maintains pre-adipocyte phenotype via modulating cytoskeletal dynamics in a manner where in the presence of pFN ECM, preadipocytes respond to insulin with proliferation and the absence results in insulin promoting differentiation (35). These data have collectively suggested an important role for FN and pFN in AT, however, no *in vivo* data has been available to evaluate this. In the current study, we provide evidence, that pFN accumulates to normal WAT. Its levels are particularly high in VAT. In SAT pFN is clearly the main FN isoform as pFN KO mice show very low detection of FN in SAT. The observation is similar to reports on other tissues where pFN was shown to contribute to nearly all detectible FN matrix, such as bone (28).

Our data from the pFN knockout shows that pFN has a most prominent role in normal AT as almost all phenotypes that we so far examined were observed on normal (control) diet-fed mice. Phenotype was only observed in male mice. pFN absence has no effect on weight gain nor on fat%, fat mass or AT depots on HFD-feeding regime. On normal diet pFN KO mouse fat% was decreased, albeit not significantly. However, significant enhancement on whole-body glucose clearance and whole-body insulin sensitivity as well as AT insulin sensitivity was detected. The whole-body effects disappeared on HFD suggesting strong signals overriding pFN function in obesity. However, the whole-body effects cannot be all attributed to the role of pFN in AT as VAT remained significantly highly insulin sensitive in the absence of pFN on HFD. The disappearance of whole-body pFN effects on HFD may arise from its role in other metabolic tissues such as liver which we did not investigate further here but show to have high pFN accrual after injections.

Our work demonstrated that pFN absence significantly promoted AT insulin sensitivity as seen in rapid *in vivo* tests and detection of pAKT engagement. The effect was significant for both SAT and VAT and the difference that disappeared upon HFD-feeding in SAT, but remained significant in obese VAT. This suggests that pFN may contribute to the SAT vs VAT homeostasis differently. The differences were also seen in the result that only in SAT pAKT phosphorylation occurred to Thr308, which may lead to activation of different pathways. The cause of the increased insulin sensitivity is likely associated mechanistically to the increased cellularity of AT in pFN which was seen in the histological analyses of the SAT and VAT, but more prominent in SAT. Vast evidence supports the concept that smaller adipocytes and preadipocytes are more insulin sensitive (7, 55) Furthermore, our RNA sequencing and qPCR data of SAT showed markers for browning, i.e., presence of beige adipocytes, which are highly insulin sensitive and may also increase the observed insulin sensitivity. Additionally, pFN ECM (or lack thereof) may also have direct effects on the cell signaling. Integrin engagement is known to boost insulin sensitivity in a manner where β1/β3 integrin activation contribute to insulin sensitivity and browning/beiging of AT (56). Furthermore, exercise-induced myokine, irisin interacts with αVβ1/β5-integrins to induce WAT browning and beige adipocyte biogenesis (57, 58). It is thus possible that the absence of pFN ECM changes other ECM constituents and/or stiffness and integrin adhesions to favor small adipocyte phenotype, preadipocyte, and beige adipocyte presence. We did not analyze BAT in this study as the accrual of pFN appeared negligible in adult mice. However, analyzing pFN levels during BAT development Our current studies will focus on dissecting these mechanisms.

Interesting changes were also seen in pFN KO VAT gene expression on HFD with downregulation of majority of the genes. Top set of these represented downregulation of skeletal development-related transcription factors, including *Grem1; Runx2, Lef1;* mineralization regulators, *Enpp1, Pitx1, Bglap3,* and osteogenesis related, *Pthr1* (parathyroid hormone 1 receptor) suggesting that adipose stem cells may have altered phenotypes in the absence of pFN. Also, tube morphogenesis was altered where fibronectins have been shown to have a major role (59, 60). However, these changes do not explain the observed increased insulin sensitivity on HFD which remains to be explored.

While the circulating pool of pFN serves as a source for the tissue ECM, our data suggested that the actual assembly is regulated locally by the presence and activity of FXIII-A transglutaminase (35, 36). pFN is a well-established substrate of FXIII-A (61–66), which is best known for its function as fibrin-stabilizing factor at the last step of the coagulation cascade (67–69). The circulating plasma FXIII-A is produced in the bone marrow by monocytes, macrophages and megakaryocytes (70, 71), however, recent advances from us and others have established its production also in tissues by several cell types (67). It is conceivable to consider that both the availability of pFN in plasma and tissue FXIII-A levels and transglutaminase activity jointly affects the pFN ECM in the tissues (35, 36, 71). We have also reported this FXIII-A-mediated assembly mechanism in mesenchymal stem cell and osteoblast cultures where pFN and FXIII-A promote osteoblastogenesis *in vitro*. *In vivo*, elimination of FXIII-A together with another TG, TG2, affects bone and marrow pFN levels, bone remodeling and mass *in vivo* (46, 72). Similarly, Malara et al. demonstrated that megakaryocyte FXIII-A promotes pFN assembly in the bone marrow (71). Interestingly, in this current study, pFN absence in SAT was associated with decreased FXIII-A levels in SAT, but not in plasma, likely because of different cellular source of the enzyme. In AT, FXIII-A is likely produced by preadipocytes, but also likely by hypertrophic adipocytes as suggested by our recent studies in humans (49, 73, 74). Thus, it is possible that pFN/FXIII-A ECM at different stages of adipogenesis may result in different effects, for example on preadipocytes/precursors in SAT and hypertrophic adipocytes in VAT.

Functions of liver and AT are tightly connected in obesity. Functional failure of AT results in exogenous triglyceride accumulation to liver (non-alcoholic fatty liver disease, NAFLD)(75). Hepatokines in turn, such as Fibroblast Growth Factor-21 and Fetuin-A, regulate AT function and insulin sensitivity. NAFLD, cirrhosis and alcoholic liver also affect pFN levels (76, 77). Our study suggests that pFN may also act as a hepatokine, and liver-metabolism mediator. In humans, data on circulating pFN levels in obese individuals, with and without NAFLD has been reported with no clear consensus (76, 78–80). pFN levels were shown higher in overweight males versus females with same body mass index (BMI) and the levels increased with age (80–82). Andersen et al. demonstrated that obesity increases FN levels in circulation and weight loss returns the levels to normal (83). Other reports documented that type 2 diabetic patients showed lower circulating pFN levels than controls of the same age groups, and that the levels showed no difference between females and males (84). In these studies it was not defined if circulating FN was the ‘classic pFN’ or cellular, EDA-FN (from tissues) which was recently linked to systemic inflammation and insulin resistance via activation of Toll-like receptor 4 (TLR4) (85, 86).

In summary, our data expands and deepens our understanding of the role of FN matrices in AT, metabolism and whole-body health and suggests a function for pFN in regulation of AT insulin sensitivity and adipocyte and adipose tissue cells pools *in vivo*. Modulation of its levels in AT may benefit overall AT health with benefits to glucose handling and improve insulin sensitivity.

## Author Contributions

M.M., M.T.K. designed study; M.M., E.M.D., M-H.N., Y.P., A.G., M.M., S.C., conducted the experimental work; M.M., E.M.D., M.T.K., L.M. analyzed and interpreted data; M.M., M.T.K., drafted manuscript; and M.T.K, L.M., edited and revised manuscript. All authors approved the final content of the manuscript.

## Acknowledgements

We wish to thank Aida Roodsari-Kasai for her technical help and funding agencies for their support.

## Funding Information

This work was supported by grants from the Canadian Institutes of Health (CIHR) PJT15089 and PJT162100 to MTK and CIHR PJT-175306 and PJT-162302, and the Alzheimer’s Association US AARG- 21-852152 to LM. MTK is a member of the FRQS Network for Oral and Bone Health and Cardiometabolic Health, Diabetes and Obesity Research Network (RSBO). EDM is supported by stipend from Le Fonds de recherche du Quebec – Santé (FRQS). MM is supported by stipend from Faculty of Dental Medicine and Oral Health Sciences and RSBO. YP received the Davis fellowship through McGill’s Faculty of Medicine and a stipend from the Canada First Research Excellence Fund, awarded to McGill University for the Healthy Brains for Healthy Lives initiative.

## Data Availability

Data is available upon request.

## Conflict of Interest Statement

The authors declare no conflicts of interest and there are no financial conflicts to disclose.

## Ethics Statement

The animal care and experimental procedures were under the Canadian Council of Animal Care and approved by the McGill University Animal Care Committee (protocol code MCGL-5188, May 3, 2023).

---

**Supplemental Figure 1.**
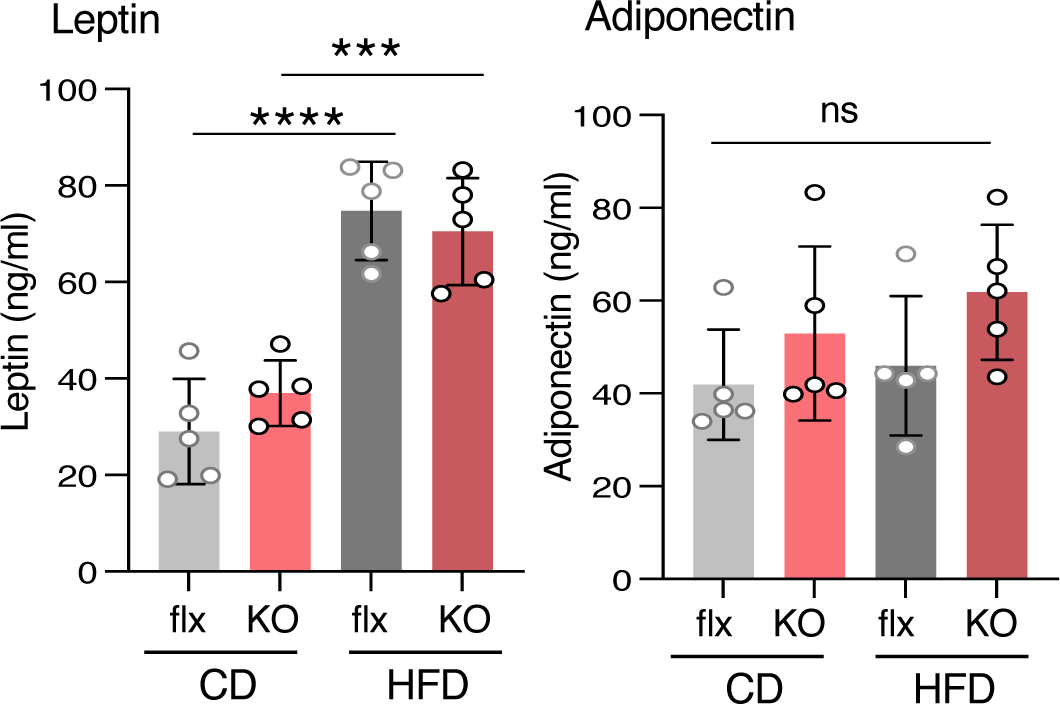
Serum leptin and adiponectin levels in pFN and flx mice. Both markers do not show significant change. Error bars represent SEM (n=5). Statistical significance is represented as: *p < 0.05, **p < 0.001, ***p < 0.0001.

**Supplemental Figure 2.**
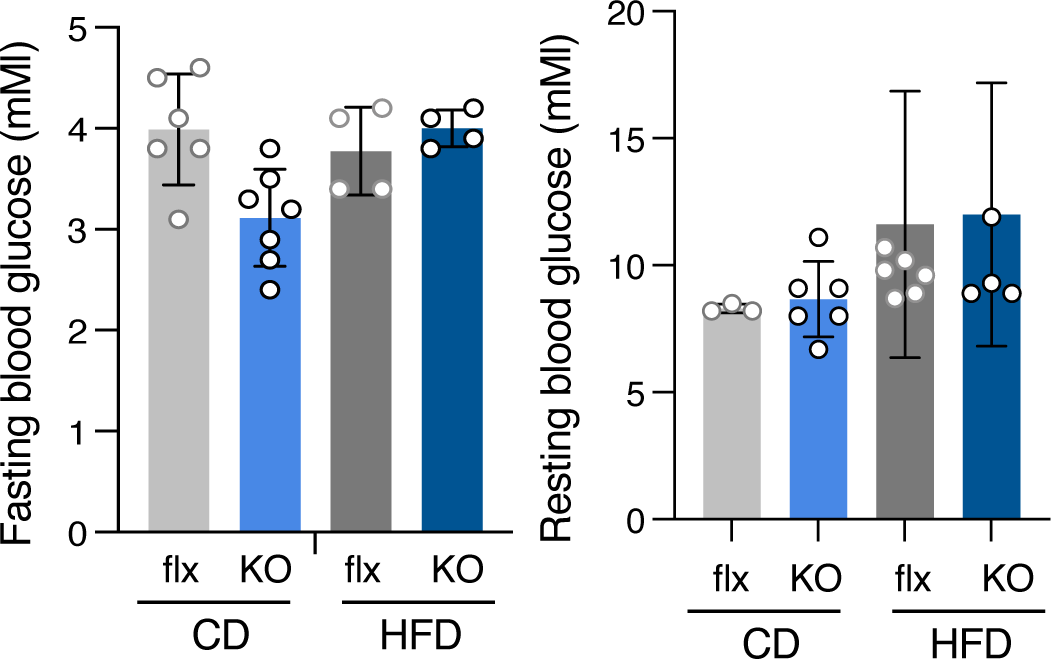
Fasting and resting glucose levels of male pFN KO. Fasting (6 h) glucose levels did not show significant differences in male or female mice on control diet (CD) or high-fat diet (HFD). Error bars represent SEM (n=5-8 per group). Statistical significance is represented as: ns; non-significant.

**Supplemental Figure 3.**
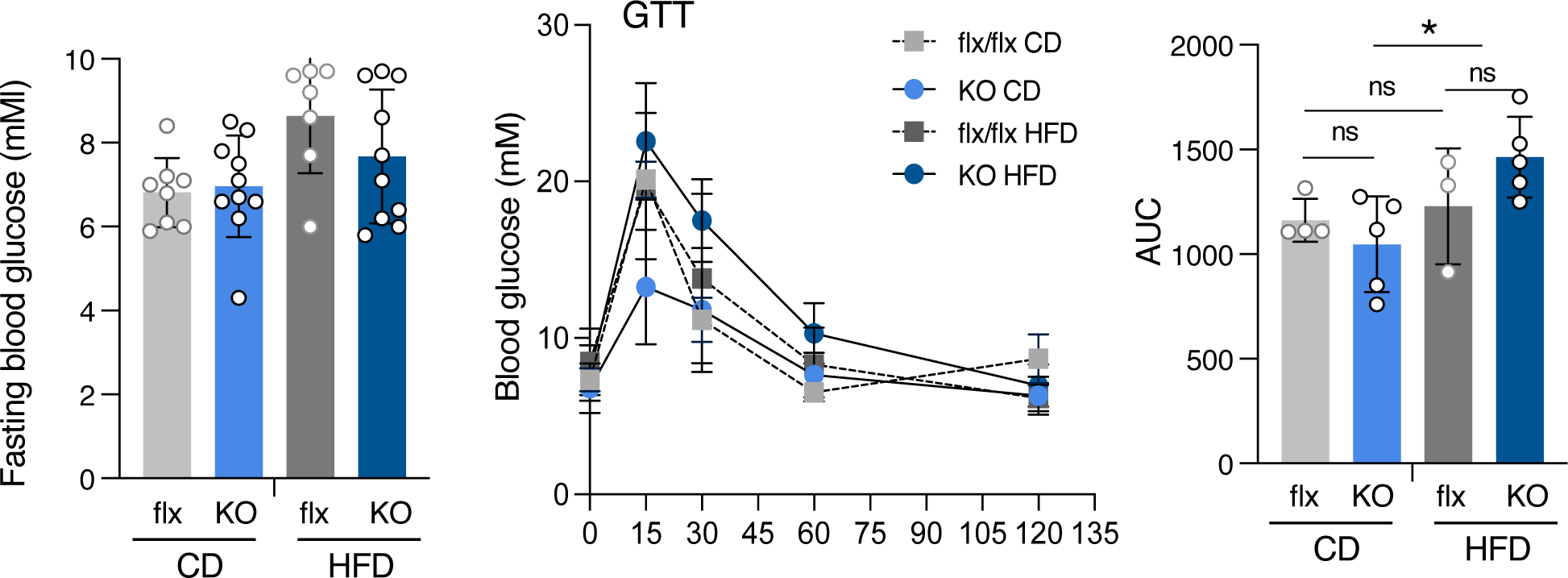
Fasting glucose levels and whole-body glucose tolerance (GTT) in female pFN knockout. Female pFN KO mice did not show significant change in glucose tolerance on either normal, control diet or HFD. Error bars represent SEM (n=5-8 per group). Statistical significance is represented as: *p < 0.05, **p < 0.001, ***p < 0.0001.

**Supplemental Figure 4.**
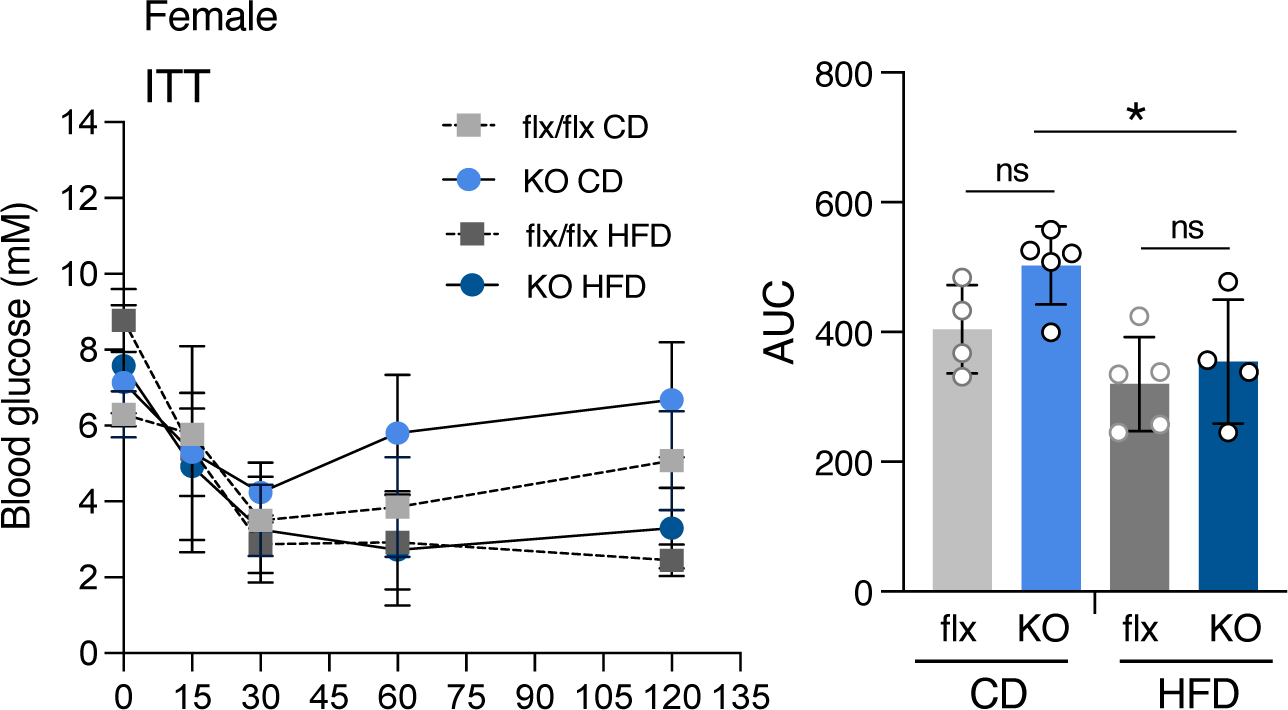
Whole-body insulin sensitivity in female pFN knockout. Female pFN KO mice did not show significant change in insulin sensitivity (insulin tolerance test; ITT) on either normal, control diet (CD) or HFD. Error bars represent SEM (n=5-8 per group). Statistical significance is represented as: *p < 0.05, **p < 0.001, ***p < 0.0001.

## Notes

### Competing Interest Statement

The authors have declared no competing interest.

